# Spatial and seasonal variation in thermal sensitivity within North American bird species

**DOI:** 10.1101/2023.03.31.535105

**Authors:** Jeremy M. Cohen, Daniel Fink, Benjamin Zuckerberg

## Abstract

Responses of wildlife to climate change are typically quantified at the species level, but physiological evidence suggests significant intraspecific variation in thermal sensitivity (non-stationarity) given adaptation to local and seasonal environments. Non-stationarity carries important implications for climate change vulnerability; for instance, sensitivity to extreme weather may increase in specific regions or seasons. Here, we leverage high-resolution observational data from eBird to understand regional and seasonal variation in thermal sensitivity for 20 bird species. Across their ranges, most birds demonstrated spatial and seasonal variation in both thermal optimum and breadth, or the temperature and range of temperatures of peak occurrence. Some birds demonstrated constant thermal optima or breadths (stationarity) while others varied according to local and current environmental conditions (non-stationarity). Across species, birds typically invested in either geographic or seasonal adaptation to climate. Intraspecific variation in thermal sensitivity is likely an important but neglected aspect of organismal responses to climate change.

## Introduction

Anthropogenic climate change is impacting wildlife at all organizational levels, from individuals to populations to species (Scheffers et al., 2016), representing a leading conservation priority for wildlife management (Abrahms et al., 2017; LeDee et al., 2021). Using traditional species distribution models or ecological niche models, ecologists typically operate at the species level to quantify responses to the thermal environment and predict the consequences of climate change (Smith et al., 2019). These approaches generally ignore adaptive capacity and phenotypic plasticity within species, implicitly assuming that thermal sensitivity, or the influence of temperature on behavior, performance, or fitness, is stationary (static) in both space and time (Jarnevich et al., 2015; Smith et al., 2019). However, emerging physiological evidence suggests that populations of a species may be locally adapted to distinct thermal conditions depending on the climate zones they inhabit, and individuals may dynamically alter their response to seasonal changes in temperature via phenotypic flexibility (Bennett et al., 2019; Louthan et al., 2021; Stager et al., 2021). Approaches assuming constant thermal sensitivity across continental spatial extents and the full annual cycle may thus be inadequate to account for the full spectrum of responses to climate change exhibited by a given species (Sultaire et al., 2022). As organisms increasingly face novel climates, understanding variation in thermal sensitivity within species will provide more detailed insights about which populations are most impacted by changing climatic conditions or extreme weather (Louthan et al., 2021; Smith et al., 2019).

Populations within a species are likely to exhibit non-stationarity in thermal sensitivity across space and time due to physiological mechanisms and constraints (Bennett et al., 2019; Louthan et al., 2021; Stager et al., 2021; Youngflesh et al., 2022; Fig. 1). Across geographic gradients, populations of wide-ranging species are likely adapted to local climatic conditions (Atkins and Travis, 2010; Stager et al., 2021). Physiological studies have suggested that within species, southern and lowland populations adapted to warm climates demonstrate warmer optimum thermal performance temperatures when compared with northern and high-elevation populations that demonstrate cooler optimums (Richardson et al., 2014; Zillig et al., 2021). Thermal breadth, or the range of tolerable conditions, is associated with the level of variability in the local climate, with ‘thermal specialists’ being found in more stable climates and ‘thermal generalists’ found in more variable climates (Bozinovic et al., 2011; Stevens, 1989). Given that climatic variability is increasing with climate change (Cai et al., 2022; La Sorte et al., 2021; Pendergrass et al., 2017), these findings highlight the importance of considering population-level variation in thermal breadth (Stager et al., 2021). In seasonal environments, non-migratory organisms must adapt to variable weather across the annual cycle via phenotypic plasticity, often undergoing behavioral and physiological changes (foraging during different times of day, seeking out refugia, gaining fat reserves, etc.) to cope with cold winter temperatures (Jimenez et al., 2020; Laplante et al., 2019). Indeed, physiological studies have revealed that organisms often fluctuate in thermal sensitivity depending on time of year (Doucette and Geiser, 2008; Hopkin et al., 2006).

**Figure 1.**
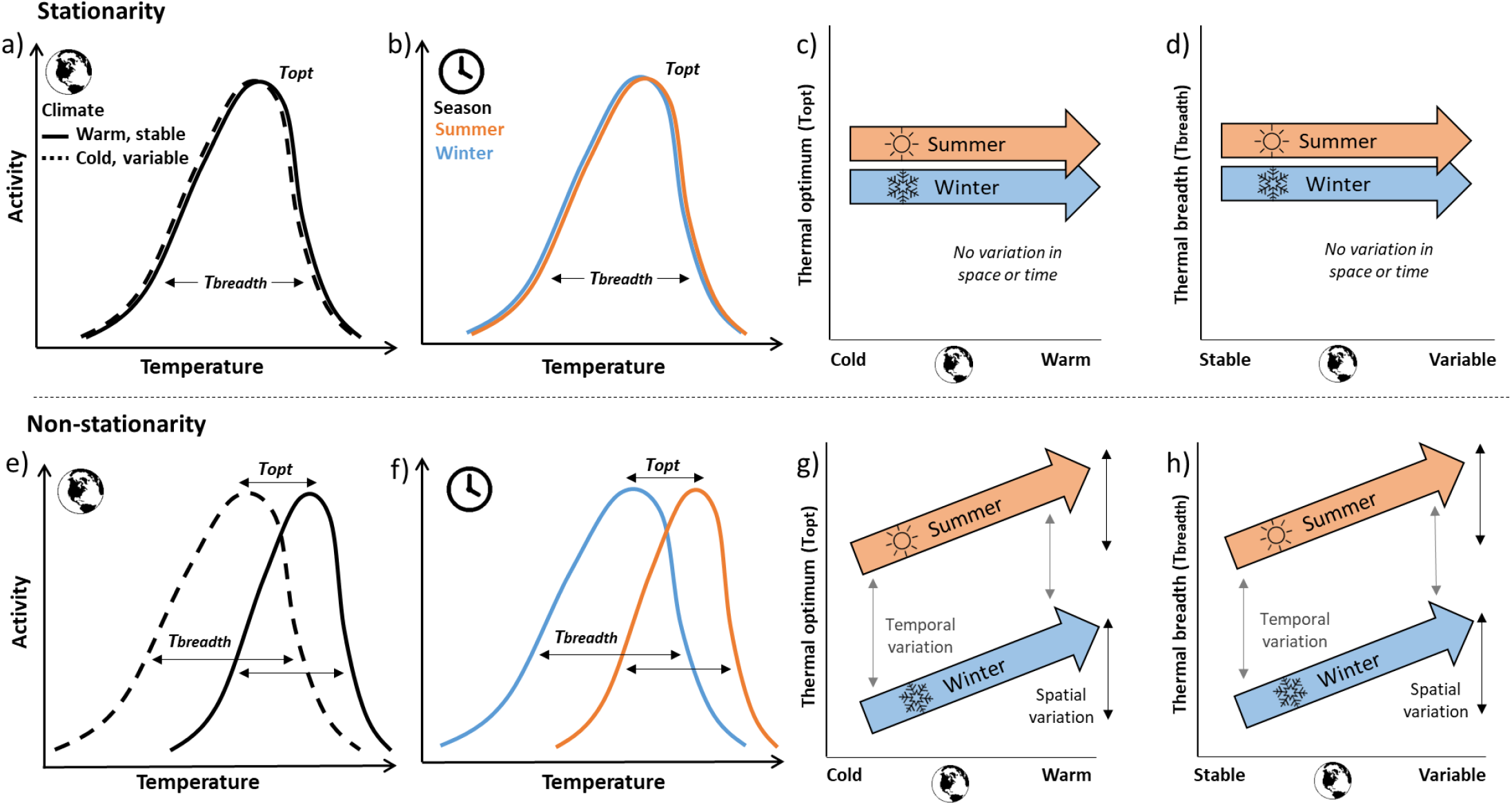
Non-stationarity in thermal sensitivity. Conceptual schematic of spatial and seasonal patterns in the thermal sensitivity of a species given **(a-d)** the assumption of stationarity, or lack of variability in thermal sensitivity within a species, and **(e-h)** non-stationarity or variation in space and time. In **(a**,**e)** curves represent hypothetical relationships between temperature and activity levels in a warm, stable climate (solid line) or cold, variable climate (dotted line). In **(b**,**f)** curves represent seasons (summer, orange; winter, blue). In right panels, variation in thermal optimum (T_opt_, temperature of peak activity; **c**,**g**) and thermal breadth (T_breadth_, range of temperatures at which activity is high; **d**,**h**) is driven by climate context only in the non-stationarity scenario. Black arrows represent the degree of spatial non-stationarity in thermal optimum or breadth, while gray arrows represent seasonal non-stationarity between seasons.

Together, evidence from physiological and behavioral studies suggests that non-stationarity in thermal sensitivity may be widespread, especially among species that occur across a wide latitudinal or elevational gradient or occupy seasonal environments (Bennett et al., 2019; Louthan et al., 2021). However, it remains unclear to what extent spatial and seasonal non-stationarity exists and whether its presence is similar across species (Louthan et al., 2021). Thus, studies are needed that can determine whether species demonstrate stationarity, meaning consistent thermal sensitivities across their ranges, or non-stationarity, where sensitivity is primarily driven by the local environment. Field- or lab-based studies conducted with one or few species (i.e., most physiological and behavioral studies) rarely capture spatial and seasonal variation in thermal sensitivity across many species occupying broad geographic regions and are unable to assess the extent to which non-stationarity is associated with certain traits across species. To evaluate the stationarity of species’ responses to their thermal environments, these relationships must be modeled at high temporal resolutions (to capture dynamic changes in temperature) across multiple seasons, and with high spatial resolution across broad geographic extents to characterize the responses of regionally distinct populations.

Across species, variation in thermal sensitivity may be mediated by morphological or life history traits (Ryding et al., 2021). For example, larger-bodied species are more common in cooler climates due to their ability to retain heat more effectively (Bergmann’s Rule; Bergmann, 1848) and larger appendages are important for effective heat dissipation for species in warmer climates (Allen’s Rule; Allen, 1877). Habitat specialists, which are more often thermal specialists than generalists (Barnagaud, Devictor et al. 2012), and species occupying forested or urban habitats, which may have more microclimates to buffer environmental conditions than open/grassland species (Jarzyna, Zuckerberg et al. 2016), may also be likely to exhibit high spatial and seasonal variation in thermal sensitivity. Thus, we hypothesize that non-stationarity is greatest in birds that 1) are small-bodied, 2) have smaller appendages, 3) are habitat specialists, and 4) occupy forested or urban habitats. Understanding which species have greater non-stationarity in thermal sensitivity – including both thermal optimum and breadth – is an important step towards anticipating organismal responses to climate change. For such species, a cold-adapted northern population may be more sensitive to warming events than a warm-adapted southern population, and a population from a stable climate may be more sensitive to increasing temperature variability than a population from a variable climate.

Here, our goal was to analyze how sensitivity to the thermal environment varies across species’ ranges and between seasons by quantifying it at high resolution across space and time. Our approach modelled the association between species occurrence rates and daily temperature and used this information to estimate population-level parameters of thermal performance. We measure thermal sensitivity as both thermal optimum, or the temperature at which a species occurs most often, and thermal breadth, or the range of temperatures at which a species occurs at 80% of its maximum rate, with daily occurrence rate as a measure of behavioral activity (Cohen et al., 2020) (Fig. 1). We used North American bird species as a case study because they are highly detectable and demonstrate strong sensitivity to weather and climate (Knudsen et al., 2011). We focused on 20 bird species from across the United States that met the following criteria: 1) broad ranges spanning latitudinal and climate zones, enabling comparisons of populations occupying diverse climates; 2) year-round presence in most of their range, enabling direct comparisons of similar populations over different seasons; and 3) ranges that overlap, minimizing variation in available thermal conditions between species that could account for differing relationships between activity levels and climatic conditions. Thus, differences in non-stationarity across species (e.g., if one species demonstrates stationarity and another with nearly the same range demonstrates non-stationarity) are a consequence of an organismal response to temperature and not simply a reflection of available conditions.

Specifically, we pose the following questions:

1. Do species vary in thermal sensitivity across their ranges (spatial non-stationarity)?
2. Do birds vary in thermal sensitivity across seasons (seasonal non-stationarity)?
3. Do species with greater spatial non-stationarity have greater seasonal non-stationarity? Species that exhibit high spatial and seasonal non-stationarity likely have increased adaptive capacity whereas a negative relationship suggests a trade-off (e.g., a species with high seasonal non-stationarity is less reliant on local adaptation).
4. Is non-stationarity mediated by species’ traits or phylogeny?

To address our questions, we present a novel analytical framework for exploring thermal sensitivity based on observational data from eBird, a citizen science initiative in which users submit bird sightings (Sullivan et al., 2014). eBird is especially useful for our approach because it has a massive data volume in the US (over 500 million records) with dense coverage, and observations are collected throughout the year, at all times of day (La Sorte et al., 2018). We leverage this dataset to identify regional and seasonal non-stationarity in thermal sensitivity for 20 species, fitting random forest models as dynamic species distribution models (SDMs) within a STEM wrapper (Fink et al., 2020, Spatio-Temporal Exploratory Models; 2010). STEM is an ensemble modeling approach that fits regional SDMs over broad spatial extents, allowing relationships between weather conditions and observations to vary spatially. We fit models using data across the full annual cycle and generated predictions for both the summer and winter seasons. In doing so, we quantified associations between species occurrence and daily temperature at local and seasonal scales to assess non-stationarity across a continental extent encompassing ∼900 million km^2^. Finally, we examined trait and phylogenetic associations with non-stationarity at the species level.

## Materials and Methods

### eBird observational data

Our overarching goal was to examine spatial and seasonal variation in the responses of North American bird species to variation in daily temperature. We compiled all ‘complete checklists’ contributed to eBird in the contiguous United States (bounding box with dimensions 25° to 47° N and 60° to 125° W) between 2004-2018. When submitting ‘complete checklists’, users indicate that all identified species were recorded, allowing the inference of non-detection for presence-absence modeling. We applied a number of filters to the data in accordance with established best practices outlined in Johnston et al. (Johnston et al. 2019). We limited checklists to “traveling” or “stationary” observations, excluding exhaustive area-counts, which are less numerous and not directly comparable with the bulk of the eBird dataset. In all checklists, subspecies information was discarded, and observations were summarized at the species level.

Likewise, we excluded checklists with extreme high values of effort (> 3 hours or > 5 km traveled, to mitigate positional uncertainty in eBird data) or extreme Checklist Calibration Index (CCI) scores (z-score < -4 or > 4), an index designed to capture inter-observer variation among eBird checklists (Johnston et al. 2019). To mitigate site selection and temporal bias, we also filtered eBird checklists by randomly selecting one observation per 5 km^2^ grid cell during each calendar week (Johnston et al. 2019). Database management was completed using *tidyverse* packages (Wickham et al., 2019).

### Distribution models

We included environmental features in each model to account for the many factors that influence species’ detection and occurrence rates. To account for variation in detection rates associated with search effort, and varying activity levels among birds at different times of the day and among observers, we included time spent birding, number of birders, whether a checklist was categorized as traveling or stationary, distance traveled, and CCI as features in species distribution models (SDMs, see below) following established best practices for modeling eBird data (Johnston et al. 2019). Further, we accounted for seasonal and daily timing by including calendar date and the time difference from solar noon in models.

To account for species preferences in landscape composition and configuration, we gathered land and water cover and topographic data corresponding to each checklist. We obtained annual landcover data from the Moderate Resolution Imaging Spectroradiometer (MODIS) Land Cover Type (MCD12Q1) Dataset, version 6 (https://lpdaac.usgs.gov/dataset_discovery/modis/modis_products_table/mcd12q1). For each checklist, we calculated the proportion of land and water classes within a neighborhood with 1.4 km radius occupied by a variety of landcover types (Hansen et al., 2000), including grasslands, croplands, mixed forests, woody savannahs, urban/built, barren, evergreen broadleaf, evergreen needle, deciduous broadleaf, deciduous needle, closed shrubland, open shrubland, herbaceous wetlands, and open savannah. Land-cover data varied annually, although we used 2017 land-cover values for checklists recorded in 2018. We also collected topographical information (median aggregations of elevation, eastness, northness, roughness, and topographic position index or TPI at a 1 km^2^ resolution) from the Global Multi-Terrain Elevation Dataset, a product of the U.S. Geological Survey and the National Geospatial-Intelligence Agency (Danielson and Gesch, 2011).

Daily mean temperatures and total daily precipitation corresponding to each checklist were compiled from Daymet, a high-resolution, interpolated grid-based product from NASA that offers daily, 1 km^2^ scale weather data across North America (Thornton et al., 2017). To account for the climate zone of each observation point, we included mean seasonal (DJF=winter, MAM=spring, JJA=summer, SON=fall) temperature and precipitation (via Worldclim; Fick and Hijmans, 2017) as additional features in random forests. The spatial resolution of our environmental features is similar to the typical radius of search effort in eBird checklists within our filters (Auer et al. *pers. comm*.).

### Species distribution models: Random Forest

The objective of the analysis was to study the relationship between species’ local occurrence rates and daily temperature for widespread, commonly detected species. We modeled responses to daily temperature in common, widespread species with sufficient data to ensure enough power to detect regional-scale variation in the relationships between temperature and occurrence across the study extent. We excluded long-distance migratory species from our analysis because winter and summer populations at the same locations are not directly comparable, although our species do move semi-locally within our spatial extent. Within the eastern or western US and Canada, we selected species with sympatric ranges to ensure that species-level differences in spatial and seasonal thermal sensitivity were not due to differences in weather availability. We divided the continent in this way to increase the similarity and overlap between species’ range extents. In the east (< 100° W), we modeled Northern cardinal (*Cardinalis cardinalis*), Blue jay (*Cyanocitta cristata*), American crow (*Corvus brachyrhynchos*), Mourning dove (*Zenaida macroura*), White-breasted nuthatch (*Sitta carolinensis*), Black-capped chickadee (*Poecile atricapillus*), Carolina chickadee (*Poecile carolinensis*), Tufted titmouse (*Baeolophus bicolor*), Carolina wren (*Thryothorus ludovicianus*), Downy woodpecker (*Dryobates pubescens*), Hairy woodpecker (*Dryobates villosus*), Red-bellied woodpecker (*Melanerpes carolinus*), and Northern mockingbird (*Mimus polyglottos*). In the west (> 100° W), we modeled Mountain chickadee (*Poecile gambeli*), Chestnut-backed chickadee (*Poecile rufescens*), Pygmy nuthatch (*Sitta pygmaea*), Bewick’s wren (*Thryomanes bewickii*), Black-billed magpie (*Pica hudsonia*), Steller’s jay (*Cyanocitta stelleri*), Anna’s hummingbird (*Calypte anna*), and Acorn woodpecker (*Melanerpes formicivorus*).

For each species, we individually fit occurrence models using Random Forests (RF; ranger package; Wright et al., 2018), a flexible machine learning method that has been used in a number of species distribution modeling problems (Mi et al., 2017) and is designed to analyze large datasets with many features, adjust automatically to complex, nonlinear relationships, and consider high-order interactions between all features. To account for spatiotemporal variation in species responses to climate across broad spatial extents, we fit RF models within a spatiotemporal exploratory models (STEM) as a wrapper (Fink et al., 2020, 2010). We used STEM to generate a randomized ensemble of partially overlapping regional models consisting of 10° x 10° cells (‘stixels’) across our spatial extent and fit independent RF models within each cell with a minimum of 20,000 checklists, producing a uniformly distributed ensemble of hundreds of partially overlapping models. Within each stixel, we assume relationships between species’ occurrence and environmental variables to be stationary. We generated spatially explicit occurrence estimates by averaging predictions from all regional RF overlapping a given location. STEM is established as an effective method for measuring non-stationary relationships between environmental features and observations (Fink et al., 2010; Johnston et al., 2015; La Sorte et al., 2017; Zuckerberg et al., 2016).

Before modeling, all data was split 75/25 into training/testing subsamples. Initial training data were further split 75/25 for model training and validation (see below). For each set, we used case-weights to equalize weighting by year, accounting for the increasing sample sizes by year generated by eBird (submissions increase 30% annually). For each model, we calibrated predicted probabilities based on a validation set calibration adjustment. Finally, we assessed the fit of each model based on a series of predictive performance metrics computed with the test data, including specificity, sensitivity, Kappa, and area under the curve (AUC).

### Partial dependence and non-stationarity metrics

To examine the regional-scale relationships between species occurrence rates and daily mean temperature, we calculated the partial dependence (Hastie et al., 2009) within each stixel. Partial dependence statistics describe how occurrence varies as a function of certain focal features, averaging across the values all other features in models (except date, see below). By averaging in this way, the partial dependence estimates capture systematic changes in occurrence associated with temperature while averaging out all other sources of variation captured by the models, including variation in detection rates and heterogeneity in search effort and among observers. For each species, we generated partial dependence estimates for both summer and winter seasons for every stixel by predicting at the median date within season (December-February dates were adjusted to a continuous scale).

We derived two measures of thermal sensitivity from partial dependence plots fit for temperature-occurrence relationships within each stixel: 1) Thermal optimum, the value of daily temperature at which predicted occurrence is maximized; 2) Thermal breadth, equal to the difference between the value of daily temperature above the thermal optimum at which predicted occurrence falls below 80% of the maximum value and the value below the thermal optimum at which occurrence falls below 80% of the maximum value. The 80% threshold is in line with many physiological studies (e.g., Angilletta Jr et al., 2002).

For both measures, we quantified the spatial and seasonal non-stationarity within each species by summarizing how thermal optimum and breadth varied across the species range and between seasons. To estimate spatial non-stationarity, we regressed mean annual temperature (bio1 from worldclim) on the thermal optimum to calculate the slope across all stixels spanning a geographic-climatic gradient within the given season, summer or winter. Similarly, we regressed mean annual temperature range (bio7) against thermal breadth to calculate the slope of thermal breadth spanning a geographic-climatic gradient within the season. A slope closer to one suggests that stixel-level thermal optimum or breadth is closely associated with local environmental conditions, while a slope closer to zero suggests that each is consistent across the species’ range. To estimate seasonal non-stationarity, we recorded the mean stixel-level difference in thermal optimum or breadth between seasons and computed a Welch’s two-sample t-test (Welch, 1938) to evaluate whether the difference in thermal optimum or breadth between winter and summer are statistically different. Greater differences suggest greater seasonal non-stationarity. Thus, we compiled six metrics of non-stationarity for each species: spatial (two seasons) and seasonal variation in thermal optimum and breadth.

All plots visualizing metrics were generated using *ggplot2* (Wickham, 2011) and *RcolorBrewer* (Neuwirth and Neuwirth, 2011).

### Influence of human observers

We explored the possible confounding influence of daily temperature on eBird observers by fitting a random forest model with daily temperature as the dependent variable and effort, CCI, landcover, topography, and mean climate features and all model parameters identical to our primary models. We then examined the explanatory power of this model, using root mean squared error (RMSE), Spearman’s rank correlation, and the partial dependency of daily temperature based on effort variables and CCI.

### Spatial predictions

We generated maps depicting spatial variation in thermal optimum throughout the range of each species across both the winter and summer seasons. First, we created a gridded dataset with 2.8 km^2^ resolution and generated model predictions of occurrence in each cell assuming 12 evenly spaced values of daily temperature ranging between 0° and 36°C, assigning a thermal optimum to each cell corresponding to the temperature at which occurrence in the cell was maximized. We held all the observation process features constant to remove variation in detectability, resulting in occurrence predictions for a standardized eBird search defined as a checklist reported by an average observer traveling 1 km over one hour. For each cell, we compiled values of land cover, elevation, and topographic features for use when generating predictions. For each species, we generated these predictions at the hour of the day when the species is most often observed based on our data, and on a day with mean annual 1970-2000 temperatures and total precipitation.

Maps were generated using the *purr* package (Wickham et al., 2019) and plotted using *RColorBrewer*.

### Species trait and phylogeny assessment

Our final goal was to determine whether spatial and seasonal variation in thermal sensitivity is associated with various avian life-history traits. We compiled information on preferred habitat (merging forest with woodland and grassland with shrubland categories), body mass (which was log-transformed) and hand-wing index from AVONET (Tobias et al., 2022). Further, we calculated species-level landcover diversity index (following Zuckerberg, Fink et al. 2016) to represent habitat generalism, based on mean partial effects of all landcover features in independent continent-wide SDMs (Cohen and Jetz *in prep*). Thus, we compiled four traits.

To assess phylogeny as a driver of non-stationarity, we calculated Blomberg’s K (Blomberg, Garland Jr et al. 2003) using an avian phylogeny (Jetz, Thomas et al. 2012) and comparing it to a null distribution of K after randomizing species’ responses 1,000 times (‘picante’ package; Kembel, Cowan et al. 2010). Finally, we fit six multivariate phylogenetic generalized least-squares (PGLS) models to assess the simultaneous influence of traits and phylogeny on each of the six non-stationarity metrics. We then fit ANOVAs to each model to assess the importance of the categorical variable (habitat preference).

## Results

Overall, species demonstrated both spatial and seasonal non-stationarity, though with considerable variation among species (Table 1; Figs. 2 & 3). During both seasons, species exhibited higher thermal optimums in warmer climates, although this relationship was stronger during winter (summer: mean β = 0.59 +/- 0.09; winter: 1.09 +/- 0.14). Birds also exhibited wider thermal breadths in more variable climates (summer: mean β = 0.1 +/- 0.04; winter: 0.09 +/- 0.05), and greater optimums (mean sample difference = 14.74 °C +/- 1.01) and narrower breadths (−2.69 °C +/- 0.43) in summer than winter.

**Figure 2.**
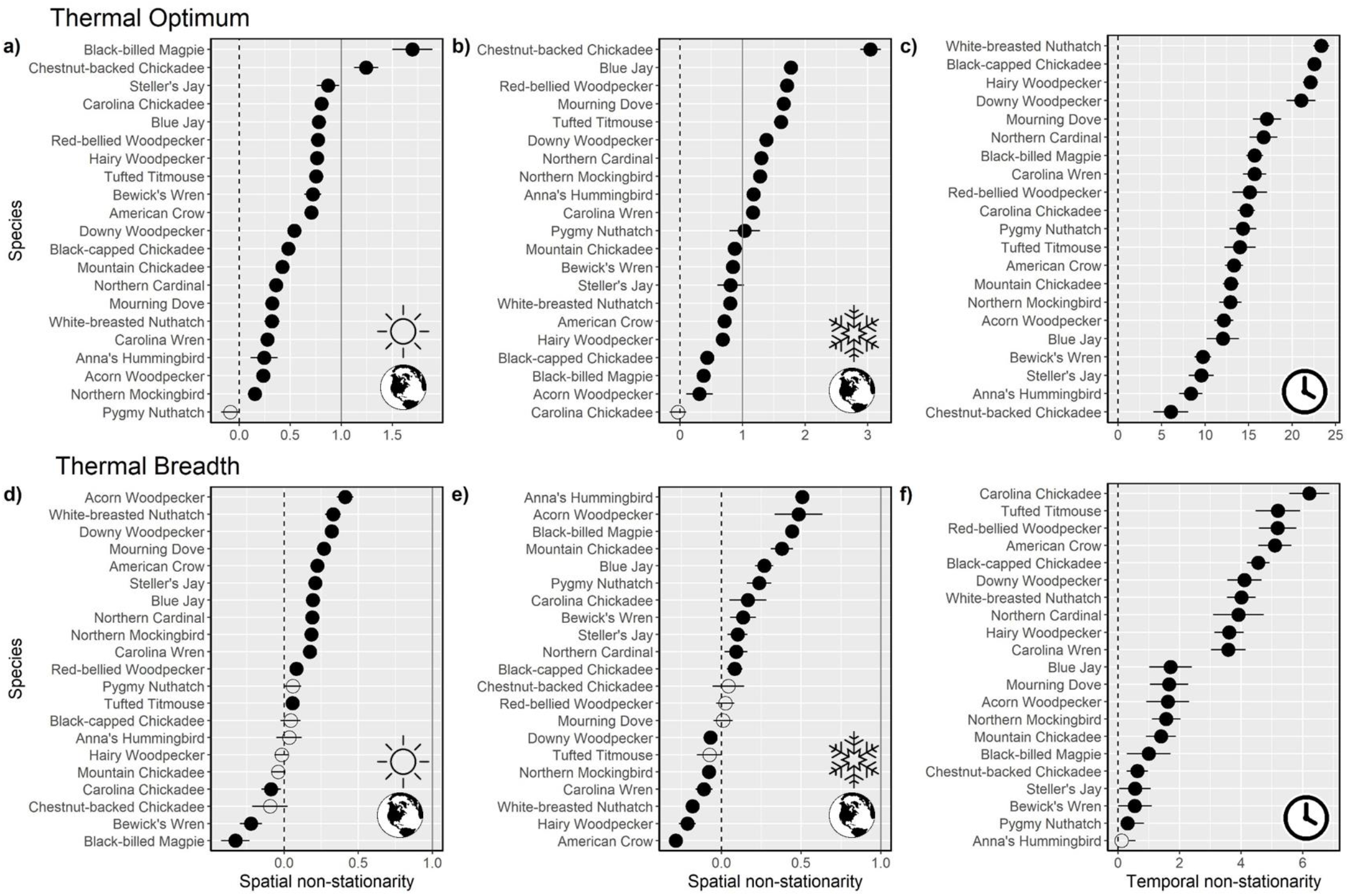
Spatial non-stationarity across 20 North American bird species. Estimates of spatial and seasonal non-stationarity in thermal optimum (a-c) and breadth (d-f). Spatial non-stationarity is defined as the slope coefficient (+/- SE) describing the regional-scale relationship between a species’ thermal optimum or breadth and the regional mean temperature or temperature range and is presented for summer (a,d) and winter (b,e) seasons. Seasonal non-stationarity is defined as the mean stixel-level difference in °C (+/- 95CI) between a species’ thermal optimum (c) or thermal breadth (f) during summer and winter seasons. The black dotted lines correspond to a value of zero, or no relationship between thermal optimum/breadth and local climate (i.e., stationarity) and gray lines correspond to one, or a 1:1 relationship, or strong spatial non-stationarity. Open circles denote species with error overlapping zero.

**Figure 3.**
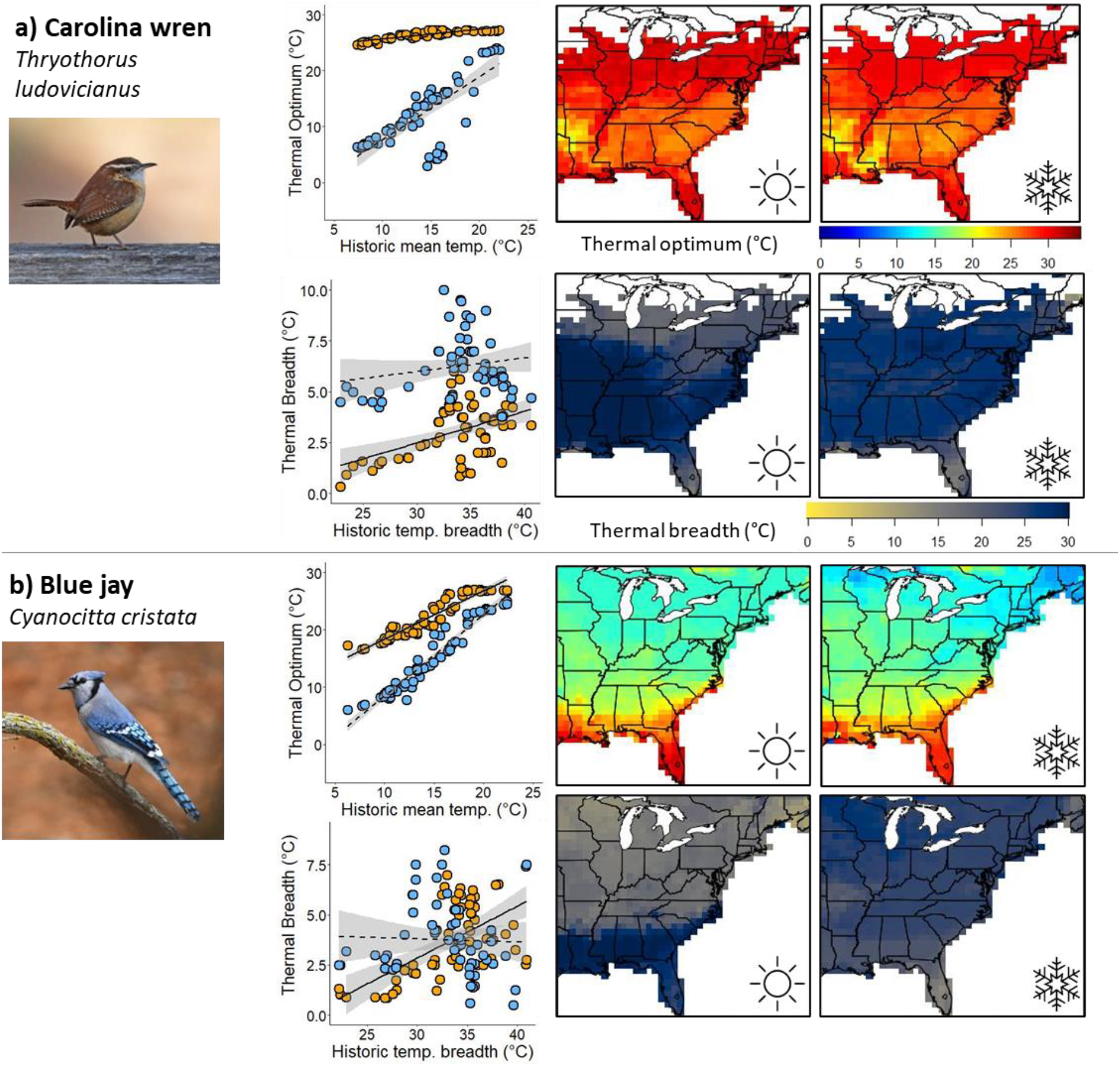
Spatial and seasonal non-stationarity differs between species. Left panels illustrate relationships between annual seasonal mean temperature and thermal optimum (the daily temperature at which activity level is greatest in each region based on model predictions; points), or relationships between historic seasonal temperature range and thermal breadth (the range of temperatures at which activity levels are above 80% of maximum) for each stixel. Patterns are given across summer (orange points, solid trendline) and winter (blue points, dashed line), with shaded 95% confidence bands. Maps visualize thermal optimums in space for each species across both seasons. (a) Carolina wren (*Thryothorus ludovicianus*) has a consistent optimum at warm temperatures with moderate spatial variation across the map, with seasonal variation in optimum occurring only in cold climates. It has moderate variation in breadth during both seasons. (b) Blue jay (*Cyanocitta cristata*) has high spatial and low seasonal variation in optimum, but more seasonal variation in breadth. Note that patterns in scatterplots may not directly correspond to those on maps because scatterplots summarize thermal sensitivity at the stixel level while maps average multiple (10-20) stixels at the point level.

In both summer and winter, all but one bird species exhibited spatial non-stationarity in thermal optimum (based on a model coefficient +/- SE not overlapping zero) across climate zones. In summer, thermal optima of two species (10%) perfectly matched that of their environment (based on a model coefficient > 1), but this increased to 11 species (55%) during winter. Spatial non-stationarity in thermal breadth was mixed, with 55% of species demonstrating shifts in winter and 60% in summer (Fig. 2). Meanwhile, seasonal non-stationarity in thermal optimum (the difference in thermal optimum between summer and winter) was observed in all birds but varied in magnitude across species, and seasonal non-stationarity in thermal breadth was observed in all species except for Pygmy nuthatch and Anna’s hummingbird (Fig. 2). Across species, we observed that birds with greater spatial non-stationarity generally had lower seasonal non-stationarity, especially in winter (optimum, β = -2.47 +/- 1.50 SE; breadth, β = -4.82 +/- 1.62; Fig. 4). We did not detect consistent effects of daily temperature on human observer effort or variation (RMSE = 8.05; Spearman’s ρ = 0.54; Fig. S1).

**Figure 4.**
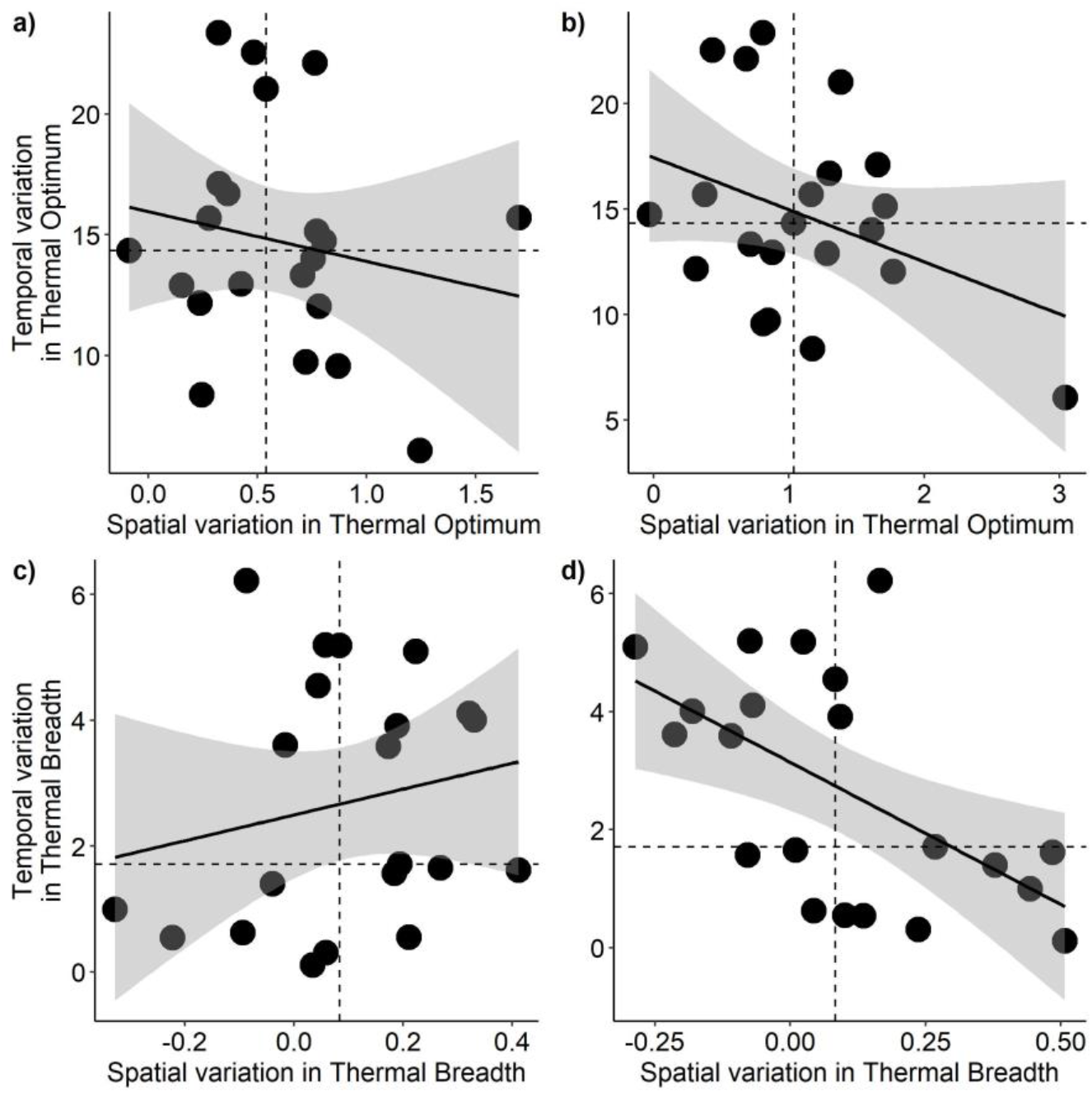
Relationships between spatial and seasonal non-stationarity in thermal sensitivity. Spatial non-stationarity (x-axes), or the slope coefficient describing the stixel-level relationship between a species’ thermal optimum or breadth and the local mean temperature or temperature range, is compared against seasonal non-stationarity (y-axes), or the mean stixel-level difference in thermal sensitivity across seasons, with points representing species. In (a-b), these comparisons are visualized for thermal optimum; in (c-d), thermal breadth. Panels (a,c) visualize trends in summer and (b,d) do so in winter. All variables were standardized to increase interpretability. Linear trendlines are given with gray shading representing 95% confidence bands. Dotted lines represent medians.

We found no evidence that phylogeny is associated with spatial or seasonal non-stationarity across species (K < 0.39, λ < 0.32, p > 0.1 for all metrics; Table S1). Most species traits were not associated with stationarity or non-stationarity either. However, habitat diversity consistently emerged as associated with spatial or seasonal non-stationarity in thermal optimum and breadth after controlling for phylogeny. For example, habitat generalists were less likely to exhibit spatial non-stationarity in thermal breadth in winter (PGLS: β = -1.70, p < 0.01), while more likely to show seasonal non-stationarity in thermal optimum (β = 36.73, p < 0.05) and thermal breadth (β = 18.19, p < 0.01; Figs. 5-6; Tables S2-3).

**Figure 5.**
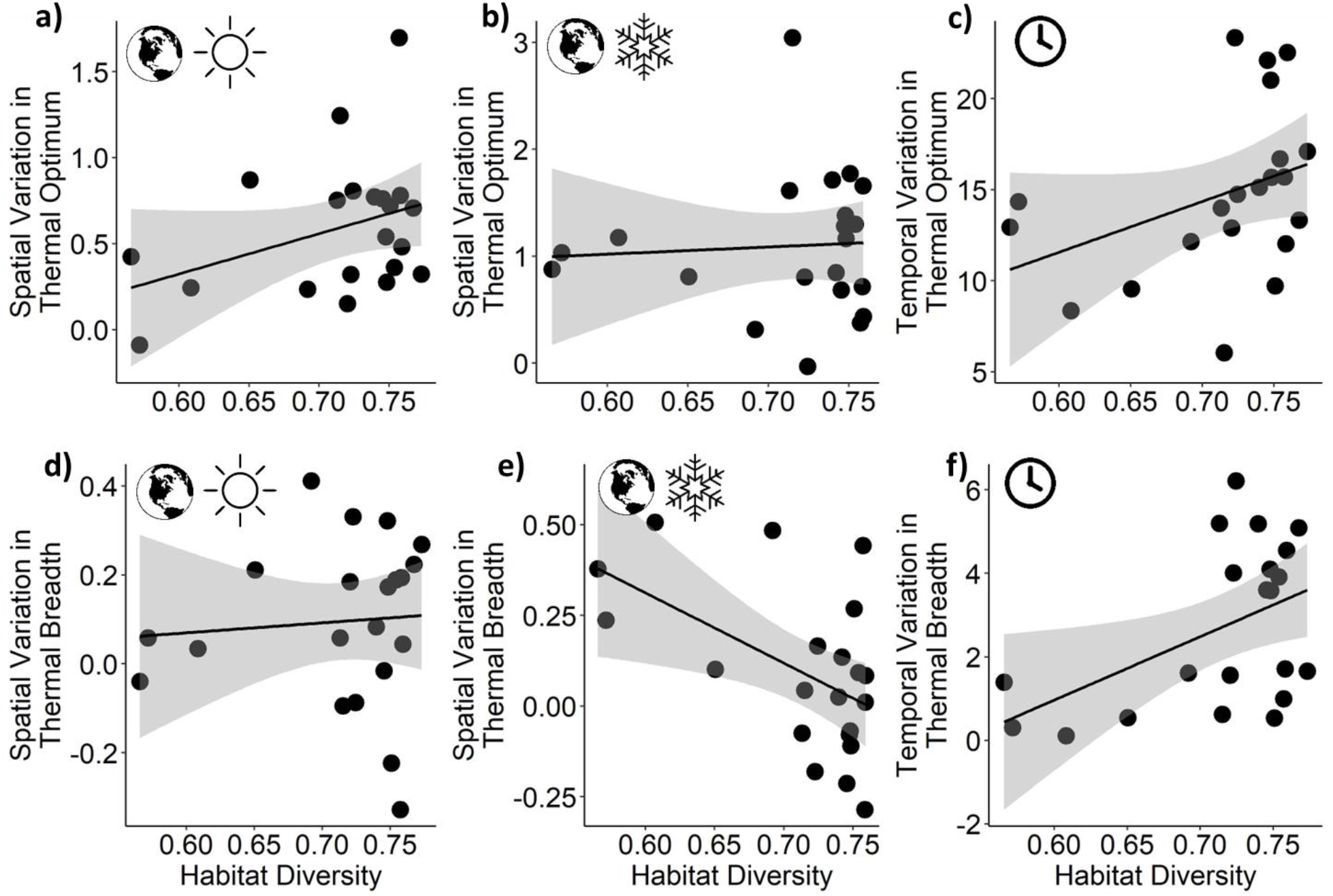
Habitat diversity is associated with the extent of non-stationarity across species. At the species level (points), partial residual plots visualize relationships between an index of habitat diversity (x-axes) and (a) spatial non-stationarity in thermal optimum in summer, (b) spatial non-stationarity in winter, or (c) seasonal non-stationarity across seasons (y-axes), based on phylogenetic least-squares models. In (d-f), equivalent relationships are presented for thermal breadth. Linear trendlines are shown with gray shading representing 95% confidence bands.

## Discussion

Thermal sensitivity and responses to climate change are typically quantified at the species level (Smith et al., 2019), but recent evidence suggests significant physiological and morphological variation among individuals below the species level (Bennett et al., 2019; Louthan et al., 2021; Stager et al., 2021; Youngflesh et al., 2022). Thus, researchers require a better understanding of variation in thermal sensitivity within species to assess when and where populations are more likely to be sensitive to weather-related effects (Louthan et al., 2021; Smith et al., 2019; Sultaire et al., 2022). However, thermal sensitivity is difficult to measure across numerous populations and multiple seasons for many species. Here, we use dynamic species distribution models that allow spatial and seasonal variation in temperature responses to identify patterns of spatial and seasonal non-stationarity in thermal sensitivity across common North American resident birds. We found that birds exhibit both stationarity and non-stationarity in responses to variation in temperature across space and time.

Our findings support recent physiological work suggesting that populations of a species vary in their thermal optimum and breadth based on geography. For both thermal optimum and breadth, most species occupied an intermediate space between complete spatial stationarity (coefficient = 0), or no variation among locations where non-stationarity that perfectly matches the local environment (coefficient = 1). Thermal optimum was more likely than thermal breadth to match local environmental conditions, with 95% of species (19 of 20) demonstrating a relationship between thermal optimum and local climate that differed from zero, and only 55% (11 of 20) demonstrating such a relationship for thermal breadth. In fact, 10 of 20 species (50%) demonstrated thermal optimums closer to one than zero, suggesting that their thermal sensitivity more closely matches the local environment than conspecifics in different regions – however, 2 of 20 species (10%) reflected a coefficient ∼1, or thermal sensitivity that matches the local environment. It has long been known that thermal breadth is highly important in terms of constraining organismal distributions, likely more so than thermal optimum (Buckley, 2010; Huey and Stevenson, 1979), and our results may suggest that thermal breadth is a more hardwired physiological constraint than thermal optimum across populations of many bird species. Across species, we found that spatial non-stationarity was infrequently associated with phylogeny or species traits, although the limited sample of 20 species limited our ability to draw broad inferences. We also found limited evidence that spatial non-stationarity in thermal breadth was greater in habitat specialists than generalists, though only during the winter season. This link was predicted because habitat and thermal generalism is often observed in the same species (Barnagaud, Devictor et al. 2012), and thermal generalists may be less likely to adapt to the local environment.

Surprisingly, all species reflected different thermal optima and 90% (18 of 20) displayed different thermal breadths across seasons, despite substantial overlap in conditions across seasons in most species’ ranges. However, this pattern may not be representative of all bird species; the species in our selection are mostly residential and thus more likely than other bird species to be seasonally flexible in thermal sensitivity. Interestingly, habitat generalism was more closely associated with seasonal non-stationarity in both thermal optimum and breadth. Therefore, habitat generalists may be selecting a strategy in which they eschew adaptation to local climates in space in favor of seasonal flexibility across the annual cycle. Finally, during winter, species with greater spatial variation in thermal sensitivity had reduced seasonal variation, suggesting a trade-off; for example, a species with seasonal non-stationarity in thermal sensitivity may not need to rely on local adaptation to climate.

Variation in thermal sensitivity across space and time may be more difficult to quantify in species that seasonally move long distances, occupy smaller ranges, or are reported less frequently, which we avoided exploring in this study. Within species that seasonally migrate long distances, seasonal variation in thermal sensitivity is difficult to measure because without knowing which sets of locations have the same individuals (e.g., information on migratory connectivity; Fuentes et al., 2022), making direct comparisons between populations over time difficult. However, recent improvements in animal tracking, even for smaller birds, will allow for direct comparisons of thermal sensitivity at the population or individual level even for migratory species (Costa-Pereira et al., 2022). Some genetic evidence suggests that populations with are northerly during the breeding season also northerly during the overwintering period (Bay et al., 2021), although this is not reliable for species which compress their ranges during winter, as do many neotropical migrants (Rushing et al., 2020). Species that occupy small ranges may exhibit little spatial variation in thermal sensitivity, as climate generally varies across large spatial scales. Although local adaptation to different climates is possible along elevational gradients, differences in data abundance between lowlands and uplands may inhibit direct comparisons between adjacent populations inhabiting each zone. Finally, assessing non-stationarity in thermal sensitivity may be more difficult for species with limited data coverage in space and time, including birds outside of North America or most other animal taxa, although citizen science observations are increasing exponentially every year (Callaghan et al., 2021).

Despite these limitations, our results provide a framework to predict how widespread, residential species with continuous data coverage may vary in population and seasonal thermal sensitivity at fine scales.

Although bird species varied in their extent of spatial and seasonal non-stationarity, it remains unclear whether non-stationarity translates to increased or decreased climate change vulnerability. Plausible explanations exist for either scenario. For example, a species exhibiting non-stationarity in space may be more vulnerable to climate change if populations are adapted to distinct thermal conditions and climates become more homogenous (e.g., northern latitudes warming faster than southern latitudes). Given non-stationarity, a continent-wide heat wave may pose a greater risk of disturbance to a northern population of a given species if it has less heat tolerance than a southern population. Alternatively, populations of a species exhibiting stationarity may be more vulnerable if southern populations already living on the edge of their thermal tolerance experience an extreme weather event, such as a heat wave. A species exhibiting seasonal stationarity may face a greater disturbance from warm weather during winter, when individuals have undergone physiological changes to suppress heat loss, than summer.

Further work should explore how variation in thermal sensitivity along a climatic gradient is related to population-level consequences to aid finer-scale conservation approaches.

## Conclusions

Researchers typically predict and measure static responses to climate change at the species level (Smith et al., 2019). In standard species distribution and niche modeling approaches, the thermal niche is treated as a static “envelope”, with climate-occurrence relationships assumed to be stationary over both species entire ranges and throughout the year (Jarnevich et al., 2015; Smith et al., 2019). Even in “dynamic” distribution modeling approaches, responses to a temporally shifting feature (e.g., weather) are assumed to be consistent across the spatial and temporal extent of the modeling domain (Milanesi et al., 2020). Further, conservationists and managers typically develop climate change vulnerability assessments and adaptation plans at species level, ignoring population-level variability. However, with the modern availability of high-resolution, high-volume, continuous observational and environmental datasets, variation in species’ responses to environmental variables, such as temperature, can now be modeled over large spatial extents and across the annual cycle to detect variation in responses to climate change as higher resolutions (Carlson et al., 2021; Latimer et al., 2018). Our results suggest that many species-level assessments of thermal sensitivity may be missing significant variability over space and time, leading to misleading climatic vulnerability assessments. Researchers must consider variation in thermal sensitivity across populations and seasons to improve understanding of climate change adaptation (Smith et al., 2019).

## Acknowledgments

We thank S. Keyser, M. Lu, and S. Sharma for their thoughtful comments on the analyses. Funding was provided by the Data Science Initiative through the Office of the Vice Chancellor for Research and Graduate Education at the University of Wisconsin. We thank the many eBird participants for their contributions and the eBird team for their support. This work was funded in part by The Leon Levy Foundation, The Wolf Creek Foundation, NASA (80NSSC19K0180), and the National Science Foundation (DBI-1939187; CCF-1522054; and computing support from CNS-1059284).

## Supplementary Materials

**Figure S1.**
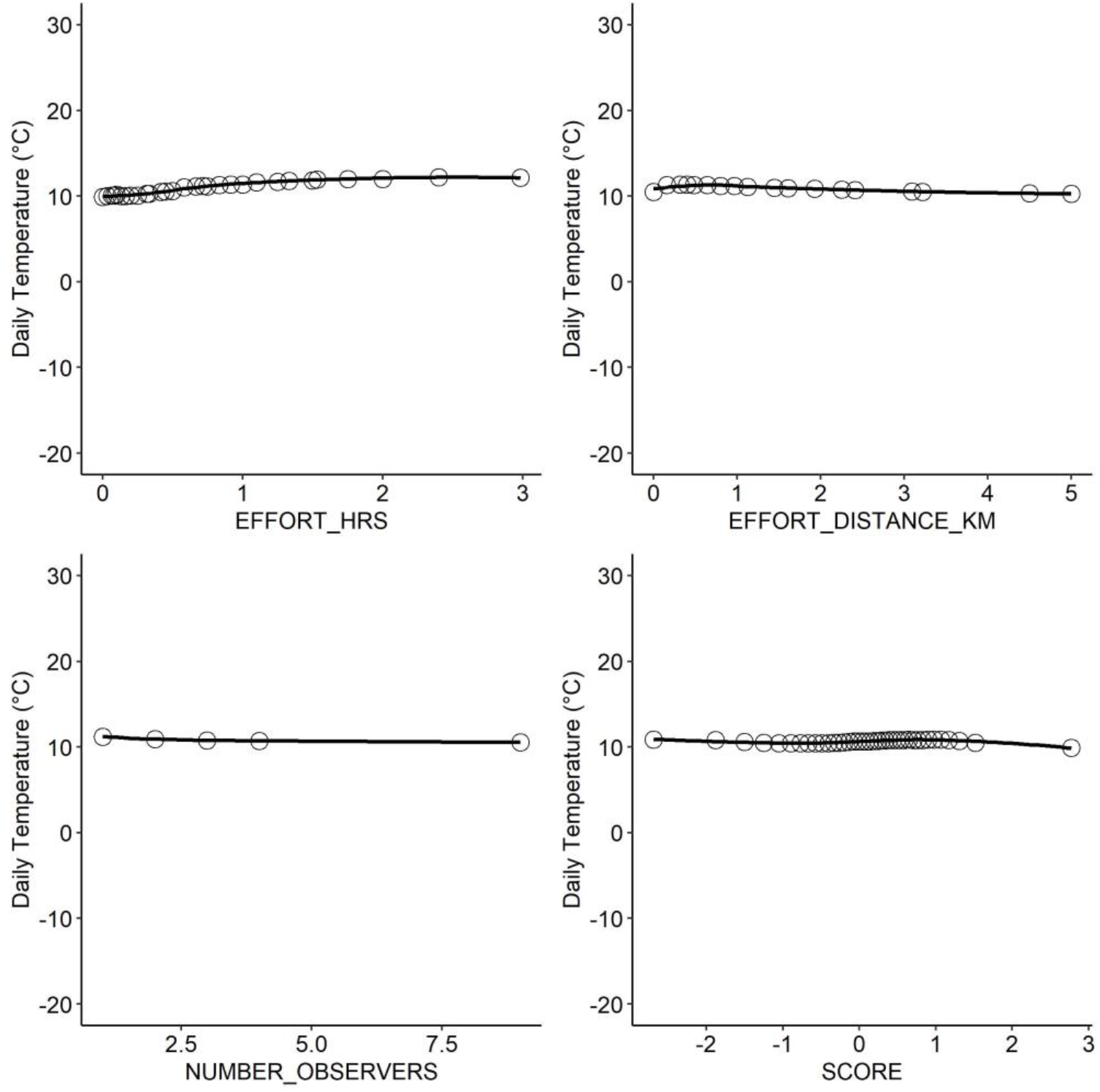
The relationship between daily temperature and human observers. Partial dependence plots based on a random forest model explicitly testing the relationship between daily temperature and metrics of human observation, including (a) number of hours birding, (b) distance traveled (km), (c) the number of observers, and (d) Checklist Calibration Index, which reflects inter-observer variation (see methods).

**Table S1.**
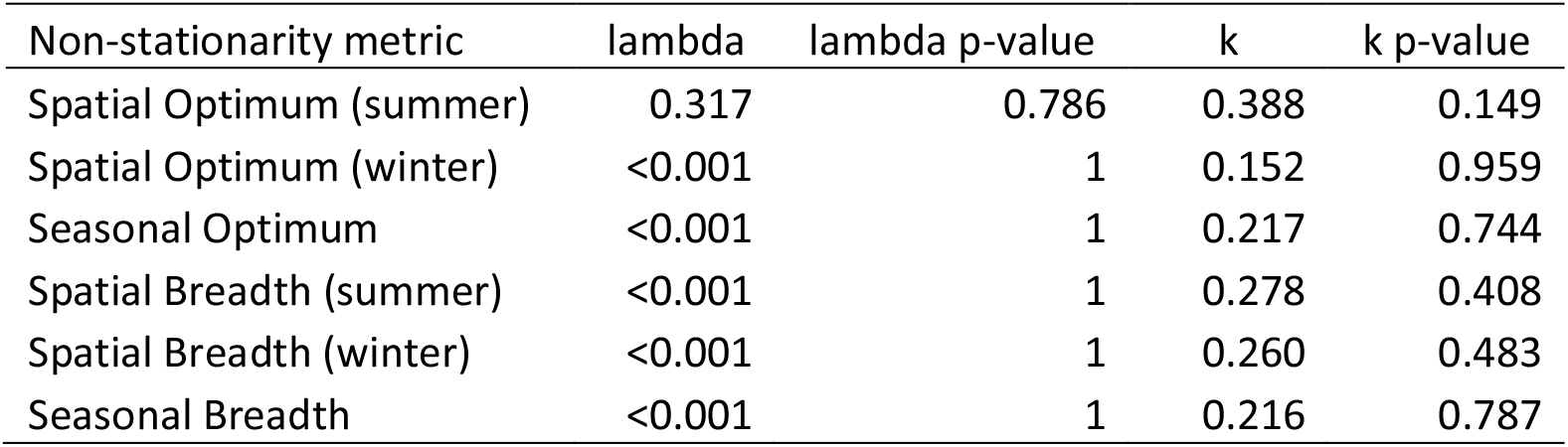
Metrics describing phylogenetic signal (Lambda and Blomberg’s k) in spatial or seasonal non-stationarity.

| Non-stationarity metric | lambda | lambda p-value | k | k p-value |
| --- | --- | --- | --- | --- |
| Spatial Optimum (summer) | 0.317 | 0.786 | 0.388 | 0.149 |
| Spatial Optimum (winter) | <0.001 | 1 | 0.152 | 0.959 |
| Seasonal Optimum | <0.001 | 1 | 0.217 | 0.744 |
| Spatial Breadth (summer) | <0.001 | 1 | 0.278 | 0.408 |
| Spatial Breadth (winter) | <0.001 | 1 | 0.260 | 0.483 |
| Seasonal Breadth | <0.001 | 1 | 0.216 | 0.787 |

**Table S2.** Summary tables from phylogenetic least-squares models associating species functional traits with spatial and seasonal non-stationarity in thermal optimum while controlling for phylogenetic structure.

| Spatial (summer) | Coefficient | SE | t-value | p-value |
| --- | --- | --- | --- | --- |
| (Intercept) | -0.939 | 1.508 | -0.623 | 0.543 |
| Body Mass | 0.064 | 0.146 | 0.439 | 0.667 |
| Habitat diversity | 1.398 | 1.037 | 1.349 | 0.197 |
| Hand-Wing Index | 0.069 | 0.373 | 0.185 | 0.856 |
| Habitat (Grassland) | 0.347 | 0.245 | 1.415 | 0.177 |
| Habitat (Human Modified) | -0.453 | 0.498 | -0.910 | 0.377 |

| Spatial (winter) | Coefficient | SE | t-value | p-value |
| --- | --- | --- | --- | --- |
| (Intercept) | 3.644 | 4.507 | 0.809 | 0.431 |
| Body Mass | -0.338 | 0.438 | -0.772 | 0.452 |
| Habitat diversity | 1.000 | 3.131 | 0.320 | 0.754 |
| Hand-Wing Index | -0.607 | 1.117 | -0.543 | 0.595 |
| Habitat (Grassland) | -0.255 | 0.734 | -0.348 | 0.733 |
| Habitat (Human Modified) | 0.667 | 1.489 | 0.448 | 0.661 |

| Seasonal | Coefficient | SE | t-value | p-value |
| --- | --- | --- | --- | --- |
| (Intercept) | -27.989 | 22.054 | -1.269 | 0.224 |
| Body Mass | 1.368 | 2.135 | 0.641 | 0.531 |
| Habitat diversity | 36.725 | 15.164 | 2.422 | **0.029** |
| Hand-Wing Index | 4.033 | 5.453 | 0.740 | 0.471 |
| Habitat (Grassland) | -3.783 | 3.583 | -1.056 | 0.308 |
| Habitat (Human Modified) | -7.227 | 7.277 | -0.993 | 0.336 |

**Table S3.** Summary tables from phylogenetic least-squares models associating species functional traits with spatial and seasonal non-stationarity in thermal breadth while controlling for phylogenetic structure.

| Spatial (summer) | Coefficient | SE | t-value | p-value |
| --- | --- | --- | --- | --- |
| (Intercept) | -0.180 | 0.769 | -0.234 | 0.818 |
| Body Mass | 0.032 | 0.074 | 0.435 | 0.670 |
| Habitat diversity | 0.232 | 0.529 | 0.439 | 0.667 |
| Hand-Wing Index | 0.024 | 0.190 | 0.127 | 0.901 |
| Habitat (Grassland) | -0.326 | 0.125 | -2.613 | 0.020 |
| Habitat (Human Modified) | 0.040 | 0.254 | 0.156 | 0.878 |

| Spatial (winter) | Coefficient | SE | t-value | p-value |
| --- | --- | --- | --- | --- |
| (Intercept) | 0.858 | 0.828 | 1.036 | 0.317 |
| Body Mass | -0.025 | 0.081 | -0.309 | 0.762 |
| Habitat diversity | -1.698 | 0.575 | -2.952 | **0.010** |
| Hand-Wing Index | 0.186 | 0.205 | 0.906 | 0.379 |
| Habitat (Grassland) | 0.212 | 0.135 | 1.571 | 0.137 |
| Habitat (Human Modified) | -0.405 | 0.274 | -1.482 | 0.159 |

| Seasonal | Coefficient | SE | t-value | p-value |
| --- | --- | --- | --- | --- |
| (Intercept) | -15.256 | 8.219 | -1.856 | 0.083 |
| Body Mass | 0.435 | 0.795 | 0.547 | 0.592 |
| Habitat diversity | 18.193 | 5.651 | 3.219 | **0.006** |
| Hand-Wing Index | 1.000 | 2.032 | 0.492 | 0.630 |
| Habitat (Grassland) | -2.411 | 1.335 | -1.806 | 0.091 |
| Habitat (Human Modified) | -0.154 | 2.712 | -0.057 | 0.955 |

